# Hexagons all the way down: Grid cells as a conformal isometric map of space

**DOI:** 10.1101/2024.02.02.578585

**Authors:** Vemund Schøyen, Constantin Bechkov, Markus Borud Pettersen, Erik Hermansen, Konstantin Holzhausen, Anders Malthe-Sørenssen, Marianne Fyhn, Mikkel Elle Lepperød

**Affiliations:** Department of Physics, University of Oslo, Norway; Department of Biosciences, University of Oslo, Norway; Simula Research Laboratory, Norway; Department of Mathematical Sciences, Norwegian University of Science and Technology, Norway

## Abstract

The brain’s ability to navigate is often attributed to spatial cells in the hippocampus and entorhinal cortex. Grid cells, found in the entorhinal cortex, are known for their hexagonal spatial activity patterns and are traditionally believed to be the neural basis for path integration. However, recent studies have cast grid cells as a distance-preserving representation. We further investigate this role in a model of grid cells based on a superposition of plane waves. In a module of such grid cells, we optimise their phases to form a conformal isometry (CI) of two-dimensional flat space. With this setup, we demonstrate that a module of at least seven grid cells can achieve a CI, with phases forming a regular hexagonal arrangement. This pattern persists when increasing the number of cells, significantly diverging from a random uniform distribution. In particular, when optimised for CI, the phase distribution becomes distinctly regular and hexagonal, offering a clear experimentally testable prediction. Moreover, grid modules encoding a CI maintain constant energy expenditure across space, providing a new perspective on the role of energy constraints in normative models of grid cells. Finally, we investigate the minimum number of grid cells required for various spatial encoding tasks, including a unique representation of space, the population activity forming a torus, and achieving a CI, where we find that all three are achieved when the module encodes a CI. Our study not only underscores the versatility of grid cells beyond path integration but also highlights the importance of geometric principles in neural representations of space.

## 1 Introduction

Since their discovery, *grid cells*, with hexagonally arranged activity in two-dimensional flat space, have been claimed to perform a range of computational tasks [1], [2]. The predominant theory posits that they play a role in path integration, a process by which an organism estimates its position by integrating self-motion information over time [3]–[6]. Several lines of experimental evidence support this theory [1], [7]–[12]. However, their exact function in the brain remains a mystery.

To unravel the functionality of grid cells, a number of models have been proposed, which can be categorised into mechanistic and normative models. Mechanistic models are typically hand-tuned such that the grid cell pattern perform path integration, while normative models are optimised allowing scientists to study the appearance of grid-like tuning curves. In this respect, normative models can have discrepancies between artificial grid-like units and biological grid cells. For instance, the grid cells in the model by Cueva *et al*. showed a square activity pattern [13]. Moreover, Sorscher *et al*. found that the grid cells in the model by Banino *et al*. had similar grid scores to low-pass filtered noise [14], [15]. Despite a significant improvement in grid score in the model by Sorscher *et al*. relative to Banino *et al*.’s model, this was only apparent in a narrow range of the hyper-parameter space of the model [16], [17]. Furthermore, Nayebi *et al*. and Schøyen *et al*. find that cells with high grid scores and randomly selected cells contribute comparably to path integration, suggesting that they might not be as crucial to the process as previously thought [18], [19]. These findings indicate that grid patterns can emerge without being directly linked to path integration, raising questions to their exact function in the brain.

To consistently generate a hexagonal grid-like pattern, other tasks or constraints that either necessitate or favour this specific solution might be at play, such as non-negativity and variations of energy constraints [13], [14], [20], [21]. However, these are not the only constraints shown to give rise to grid-like solutions. Recently, Xu *et al*. and Gao *et al*. introduced a conformal isometry (CI) constraint [21], [22]. In geometric terms, a conformally isometric neural representation preserves the angles and distances of the original space. Subsequently, others have shown that similar constraints tend to yield grid-like spatial representations [23], [24]. This could suggest that the spatial representations of grid cells indeed adhere to these geometric principles.

Previous studies also suggest that grid cell representations form a metric for space [1], [15], [25]–[30]. This is inherently realised if these cells indeed form a conformal isometry of space. This implies that distances represented in the grid code would correspond proportionally to distances in space. In the opposite case, an animal could perceive flat space as uneven and, hence, shortest paths (homing vectors) as non-straight paths.

How, then, can we explore whether grid cells represent a conformal isometric map of space, and what type of model should be used? Normative models typically have fewer assumptions on the functional form of the emergent cell types compared to mechanistic models. However, the added degrees of freedom allow a variety of spatially tuned cells to emerge, including heterogeneous gridlike cells [18]. This complexity obscures the association between cell types and function. For a given set of heterogeneous grid cells, for example, it could be hard to determine which conditions lead to a conformal isometry. Instead, we consider a simplified model with stronger assumptions on the functional form of the grid cells proposed by Solstad *et al*. [31].

In the model by Solstad *et al*., grid patterns are generated by superposition of plane waves and determined by three free parameters: scale, orientation and phase. While orientation and scale are known to be consistent within grid modules [32], and other models suggest geometric progression in scale between modules [33] and a specific relative orientation [24], the distribution of grid phases is often assumed to be uniform. Experimentally, Yoon *et al*. also suggests uniformity, as is implied in most continuous attractor grid cell models (and derived in most adaptation models in regular geometrical environments) [3], [34]. However, it has also been indicated in experimental data that the spatial phases may deviate from a random uniform arrangement [35]. We, therefore, consider the arrangement of phases as the free parameter for making a module of grid cells a conformal isometry.

In our computational experiments, we find that a module of at least seven grid cells can constitute a CI of two-dimensional flat space. In this case, the phases arrange as the edges and center of a regular hexagon. When optimising many grid cells for a conformal isometry, we also find that the phases do not follow a random uniform arrangement, but are instead significantly regular. Further investigation reveals that the distribution of the phases are strikingly hexagonal. This finding is not only a theoretical insight but also provides an experimentally testable prediction.

We also find how isometry scale, energy and resolution vary with cell count and firing rate. Of particular interest to the normative modelling of grid cells is the realisation that a module of grid cells has constant energy expenditure everywhere in space when its representation is a CI. This provides an explanation for why energy constraints are important when generating grid-like solutions.

Lastly, we investigate other objectives, such as generating a persistent torus in neural representation and the feasibility of linear decoding of 2-D spatial position from grid cell modules. Our findings indicate that at least six grid cells are needed to encode a torus, and that Cartesian coordinates cannot be perfectly linearly decoded from a grid cell module. This has implications for normative models of grid cells, suggesting that models with a linear decoding layer might not be suitable for decoding Cartesian coordinates from a pure grid-like, upstream layer.

## 2 Results

### 2.1 Grid cells as a spatial representation

We begin by introducing the model generating hexagonal patters to provide intuition about grid phases. We adopt the model from Solstad *et al*. (described in Grid cell model), representing idealised grid cells as a superposition of three plane waves (refer to Fig. 1a). This configuration results in a grid cell forming a periodic pattern across 2-D planar space. The smallest repeating segment of this pattern, termed a *unit cell*, forms a hexagon (illustrated in Fig. 1b). Given that all cells within a module share the same scale and orientation, with only their phases varying, a population vector of these grid cells maintains periodicity with the unit cell (as depicted in Fig. 1c).

**Figure 1.**
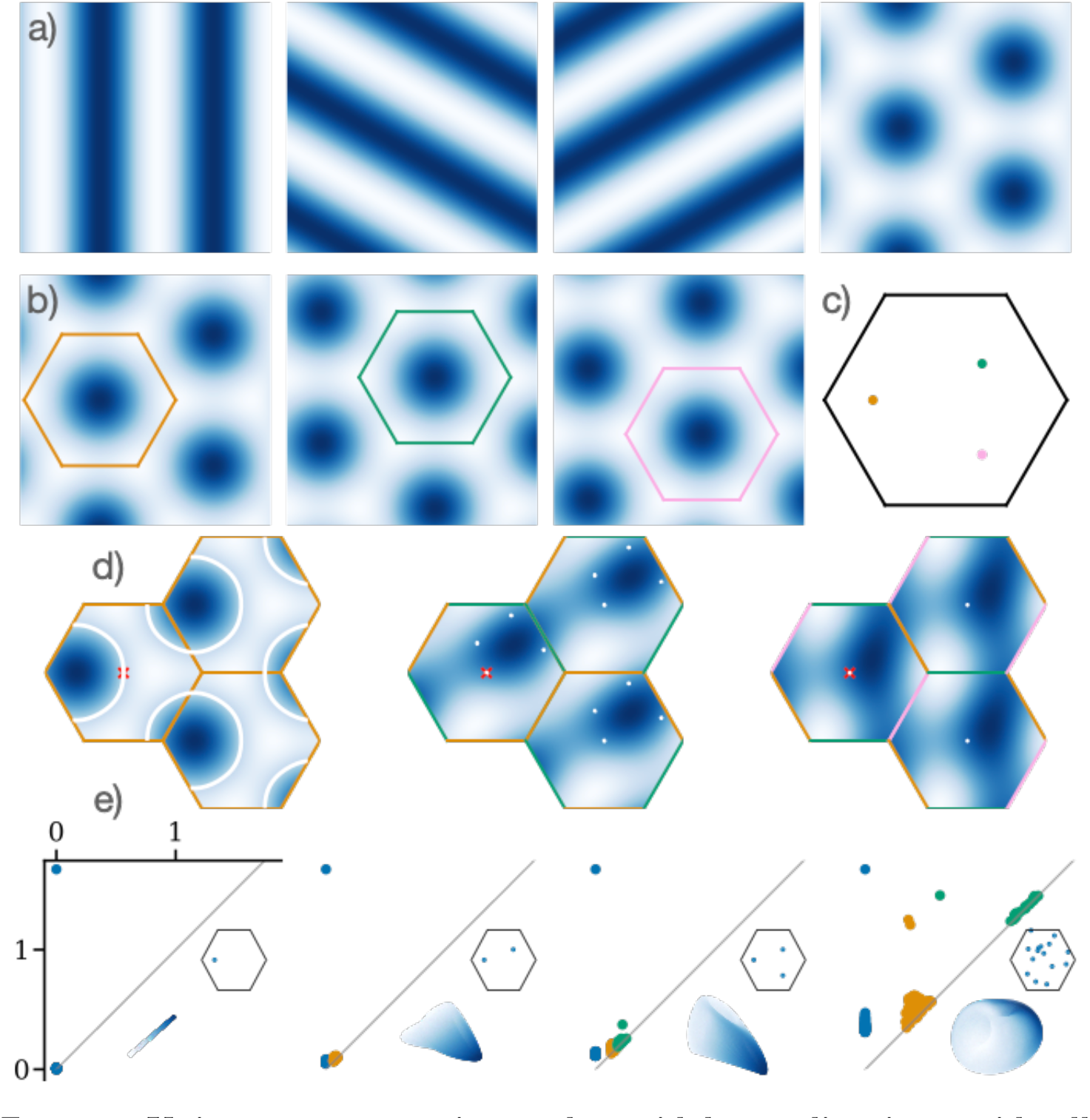
Unique representation and toroidal encoding in a grid-cell module with varying phases. **a)** Illustration of grid cells modelled as a superposition of three plane waves. **b)** Visualization of three distinct grid cells from the same module, each with their respective unit cells superimposed, demonstrating the periodic patterns. **c)** Depiction of the phases for the three grid cells in b) within a shared unit cell. **d)** Population vector correlation of 1, 2 and 3 cells’ activity (from left to right) relative to the activity at the red cross. White lines and dots highlight ambiguously represented points. For 1 and 2 cells, ambiguities arise within the unit cell. **e)** Collection of persistence diagrams, low-dimensional projections, and phase arrangements for four different grid-cell modules, varying in cell count and phase. The sequence demonstrates that modules with fewer cells (1-3) primarily encode a single persistent connected component, whereas a module with a larger number of cells (shown here with 15) successfully encodes a persistent torus.

Our first investigation focuses on the conditions necessary for a module of grid cells to represent its unit cell uniquely. Specifically, this means identifying the requisite number of grid cells and their specific phases to establish an injective map (a function which maps every position in space to unique neural population states). In theory, a set of three grid cells can uniquely represent its unit cell (proof provided in Theorem A.1). This concept is visually represented in Figure 1d), where the colours indicate the correlation between population activities at various spatial locations compared to a fixed point (marked by the red cross). The white contour lines highlight regions where the population activity is identical to the red cross location. For populations of one and two grid cells, there are ambiguities within the unit cell. However, these ambiguities can be resolved in a population of three grid cells.

Our second line of inquiry examines the conditions under which the population activity of a grid-cell module forms a toroidal manifold, a concept supported by both theoretical [25], [36] and recent experimental studies [37]. While three grid cells can uniquely represent space within the unit cell, this configuration falls short of having the structure of a torus. Prior research has suggested that approximately 20 grid cells with randomly arranged phases are required to form a torus [38]. Our observations align with this, as seen in the persistence diagram in Fig. 1e). However, we find that a torus can be formed with as few as six cells with an optimal phase arrangement (see Fig. A3).

### 2.2 Why phases matter

As previously stated, we argue that a fundamental attribute of a module of grid cells should be its ability to represent spatial distances equally everywhere. This concept aligns with the idea of encoding space as a conformal isometry, ensuring equal representation in all directions. To illustrate the importance of this, consider the opposite unfair scenario: a football field where one side appears disproportionately larger than the other (see Fig. 2a)). The metric tensor, a mathematical structure on a manifold (such as a surface) that allows defining distances and angles, can be used to describe how a representation distorts the variable(s) it encodes. Following the typical experimental setup, we restrict ourselves to the case of encoding two-dimensional Euclidean space by a module of grid cells. In this case, the metric tensor is a matrix *G ∈* ℝ^2*×*2^ at each point on an intrinsically two-dimensional manifold. Its components *G*_*xx*_, *G*_*yy*_, and *G*_*xy*_ = *G*_*yx*_ (illustrated in Fig. 2b)) describe distortions in canonical and joint directions. To represent spatial distances equally everywhere is to attain a metric tensor that is diagonal, i.e., *G* (*<u>r</u>*) = *σI*, where *I* is the identity matrix and *σ* is some scalar.

**Figure 2.**
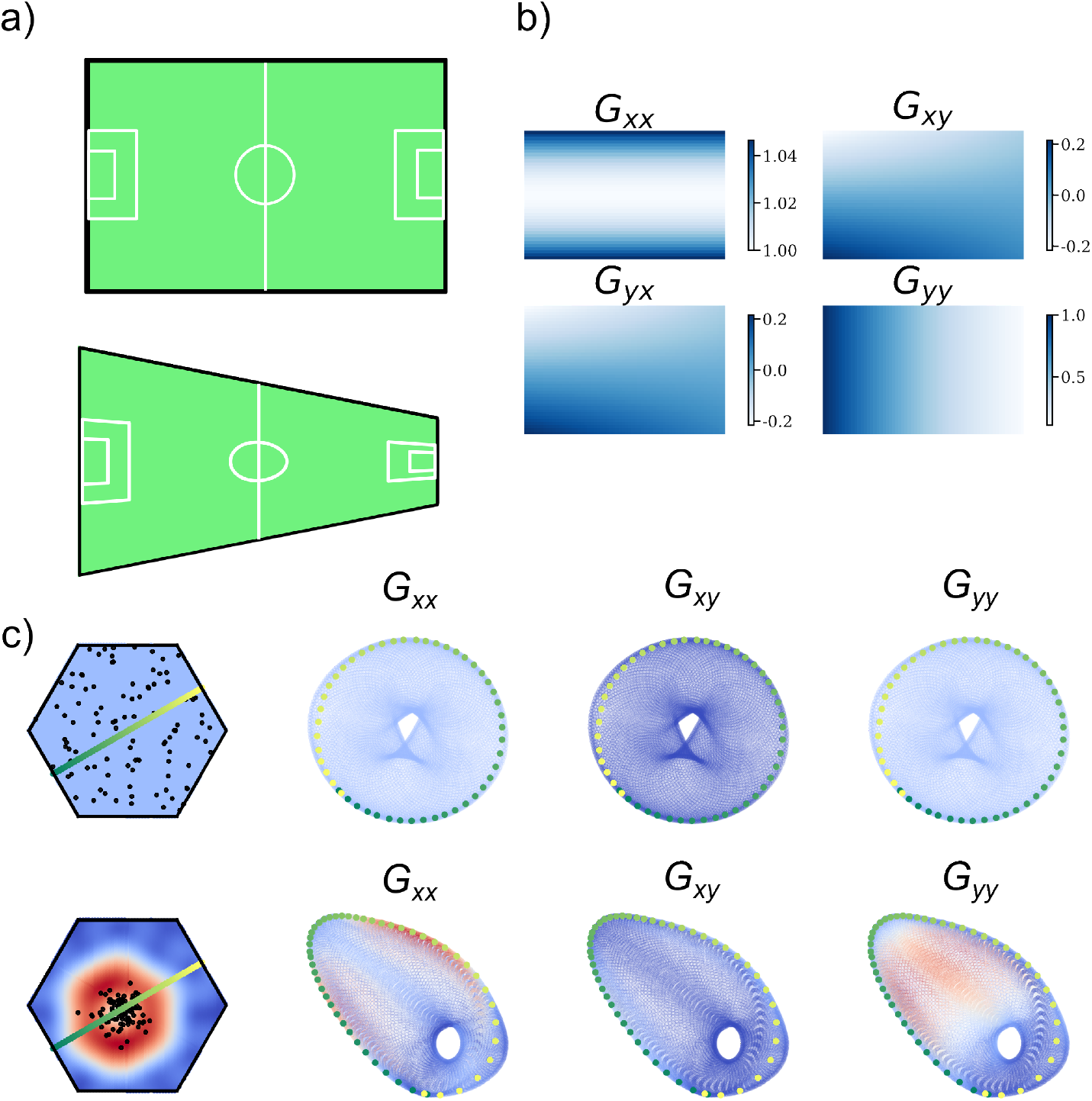
Distortions in spatial representation are encoded in the metric tensor. **a)**: An illustrative example of an ordinary football field (top) and a distorted version (bottom), in which vertical distances shrink as one goes to the right. **b)** The metric tensor components show how distances in **a**) are locally deformed in each direction. For example, the *G*_*yy*_ component shows that as one goes to the right, distances in the vertical direction will shrink, whereas the *G*_*xx*_ component shows that as one goes up or down, distances to the right will expand. **c)** Phase samples of 100 grid cells inside a unit cell (upper left in each set of four figures), along with a three-dimensional UMAP projection of the generated activity coloured by the metric tensor components (indicated by labels). Unit cell plots are coloured by the determinant of the metric tensor. The example on the first row shows a low-dimensional projection of a conformally isometric torus. In this case, the metric tensor is flat, and its trajectories are not distorted. The example on the second row shows a projection of the torus, when the phases are sampled densely around a small region. In this case, trajectories crossing phase-dense regions appear more dense on the manifold.

In the model of a grid-cell module, the phases are the only free parameters. We therefore study how the phase arrangement influences spatial representation as measured by the metric tensor. We start by considering trajectories on the neural manifold. For an optimal set of phases (as defined in the subsequent subsection), each trajectory in the environment will be similarly represented as trajectories along the internal representation. For example, a module with densely clustered phases would expand its internal representation in phase-dense areas and contract it in sparse regions. Both effects are visually demonstrated in Fig. 2c).

### 2.3 Optimal phase arrangements

Equipped with an understanding of the metric tensor and an intuition of how phases impact grid cell representations of space, we begin our research. We explore the minimal number of grid cells required for forming a conformal isometric map of space and the implications of increasing the number of cells in a module. Subsequently, we investigate the phase arrangements for both minimal and large modules.

We start by optimising the phases of a module of 1-14 cells (see Fig. A1) for a conformal isometry using the loss function detailed in Equation 4. We find that there is no appreciable decrease in loss for less than seven cells. For seven cells, however, the loss drops twelve orders of magnitude from *≈* 10^1^ to *≈* 10^*−*11^ during training (optimisation details are described in Training details). This considerable drop in loss indicates that a minimum of seven grid cells is necessary to form a conformal isometry. For 8-14 cells, the loss is also strongly reduced, although this reduction varies with cell count.

We seek to further understand *why* seven is the minimal number of cells required for forming a CI. We find that the phases of the optimised seven-cell module are consistently arranged as the corners and centre of a regular hexagon, as shown in Fig. 3a). Seven cells forming a CI concurrently form a torus, as evidenced in Fig. 3b) and c). The associated Voronoi diagram shown in Fig. 3d) reveals a partitioning of space into similar-sized hexagonal regions centred at each phase. This suggests that cells are optimally arranged to equally encode a single portion of the underlying stimulus space (here taken to be the hexagonal domain in physical space), similar to the coding strategy proposed in [39]. This can be understood from convex code theory for neural coding [40], [41], where each neuron has a receptive field, i.e. a convex region of stimulus space to which it responds. Karoubi *et al*. prove that a *good cover* of a toroidal stimulus space, requires a minimum of seven cells [42], meaning that the overlap of any fields is contractible, disallowing two fields to intersect at disjoint places. Such an encoding agrees with the map-colour theorem, stating that a subdivision of a torus into arbitrarily shaped regions requires at least seven colours to tile the surface so that no two adjacent regions have the same colour [43].

We further explore whether the hexagonal phase arrangement is arbitrary or represents a unique and robust solution. To test this, we conduct a series of grid searches to probe for solution invariances. Initially, we manipulate the phases as if they were positioned at the vertices and centre of a hexagon, varying their orientation and radius and find that the solution is unique to these parameters (see Fig. 3f). We find the radius of the hexagonal arrangement of the phases (coloured orange in Figure 3a) to be 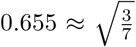 and rotated by 10.9 degrees relative to the unit cell radius. This is in line with theoretical values of the 7-colour map tiling as shown in Figure A2. Furthermore, we observe that a common phase translation does not alter the loss, suggesting translational invariance of the solution (see Fig. 3g). We confirm analytically that solutions are invariant to translations (see Independence of CI solutions).

To delve deeper into the interdependencies of phase arrangements, we experiment by translating a single phase while maintaining the others fixed. This approach highlights the collective role of phases for achieving a conformal isometry, as shown in Fig. 3h). Interestingly, we find analytically that conformal isometry solutions are independent, allowing for the combination of multiple solutions to form new, valid solutions (see Independence of CI solutions). This solution independence is computationally verified by generating a module consisting of 15 uniform random phase-shifted copies of the hexagonal solution from Fig. 3a), with the resulting phases and kernel density estimates (KDEs) depicted in Fig. 3k). Notably, the conformal isometry loss scales linearly with the number of modules copied, reinforcing the notion of solution independence.

These analyses collectively affirm that the seven-cell phase arrangement forming a regular hexagon is not a product of chance but a genuine optimal solution. However, it is important to note that while the specific arrangement of phases is critical, the overall position of the hexagonal solution is subject to arbitrary common phase shifts, indicating a form of positional invariance. Moreover, the independence of solutions suggests a wide landscape of conformal isometry solutions, potentially available for any combination of “prime”solutions, such as factors of seven cells.

Another interesting aspect is the energy expenditure of the cell ensemble. Ideally, it should be constant across all spatial locations, avoiding preferential treatment of any particular area. Interestingly, we observe that the energy (the L2 norm of the activity) of a grid module optimised for conformal isometry is constant everywhere in space (see Fig. 3e)). We can further interpret this geometrically, that points with a constant norm live on an N-sphere. Combining this observation with our knowledge of the grid module also encoding a persistent torus, we can infer that the torus lives on a 3-sphere, or *T* ^2^ *⊂ S*^3^ *⊂* ℝ^*N*^, where *N* is the number of neurons. This shows an interesting relationship between constant energy, toroidal topology and conformal isometry - they can all be achieved simultaneously with a module of grid cells, and it can be enough to optimise for conformal isometry to achieve all three.

Finally, we were curious whether a conformal isometry could still be learned, and how the phases would arrange when including more cells. We trained 50 models with 100 cells from a random uniform initialisation, each for 500 training iterations. The loss starts at *≈*10^0^ and drops to around 10^*−*7^ with a noticeable saturation at around 180 training steps. This stark drop to a vanishing small loss shows that the model has learned a conformal isometry.

To explore the resulting phase arrangement, we employ Ripley’s H-function, a spatial statistic that enables us to compare the regularity of optimised phases to a random uniform distribution [44]–[46]. We find that the optimised phases are significantly dispersed (see Fig. 3i)). Given the observed regularity in the phase arrangements of both many cells and the specific hexagonal formation of seven cells, we hypothesised that a larger assembly of cells might also exhibit a hexagonal phase distribution. By inferring the phase distribution using kernel density estimation and assessing its grid score across various kernel bandwidths, we confirmed that the optimised phases indeed form a hexagonal pattern (see Fig. 3j)). Fig. 3l) shows the optimised phases, overlaid with a kernel density estimate for different bandwidths, alongside their corresponding autocorrelation. When compared to a random uniform phase distribution (Fig. 3m)), it is evident that the phases of grid cells, when optimised for conformal isometry, are not randomly scattered but are distinctly regular and hexagonally arranged. This finding not only supports our hypothesis but also underscores the non-random, structured nature of phase distributions in grid-cell modules optimised for conformal isometry.

### 2.4 Benefits of many cells

Although three grid cells can encode a bijective transform, six are necessary for forming a torus, and seven cells are needed for a conformal isometry, more cells may carry important computational benefits. A larger ensemble of cells not only reduces the effects of cell death, but can also increase robustness and fidelity in the spatial representation. This section studies how increasing the cell count impacts the volume of the encoding manifold. Specifically, when the model adheres to a conformal isometry, this volume can be quantified using the *conformal scale*. We investigate how this affects energy consumption and the resolution of the spatial representation.

We initiate our exploration with a one-dimensional example, considering the firing rate of *N* cells encoding a line interval *r ∈* [0, 1] as

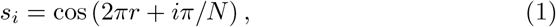

where *i* is the cell index. Fig. 4a) illustrates the tuning curves for populations of 2, 3, and 10 cells, revealing how these configurations collectively form a ring within an *N* -dimensional cube (Fig. 4b)). This gives a locally conformal map between the external line and the internal ring representation. The ring’s size, or radius, expands as 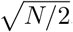.

Figure 3. : **i)** Ripley’s H function comparison between learned (blue) and random (orange) phases, with significant differences marked by stars (*p* 0.01) as determined by a permutation test. **j)** Grid score analysis of learned versus random phase distributions, inferred through Gaussian kernel density estimation, plotted against varying kernel bandwidths. **k)** Visualization of 15 replications of the phase solution from a), each subjected to random (common) uniform phase shifts, displayed alongside their kernel density estimates (bandwidth=0.2), with the grid score indicated in the title. **l)** (top) Learned and inferred phase distributions, with phases superimposed. Titles indicate the KDE grid score. (bottom) Autocorrelation of corresponding KDEs. **m)** Similar plots as in l) but for random phase distributions, providing a comparative baseline.

**Figure 3.**
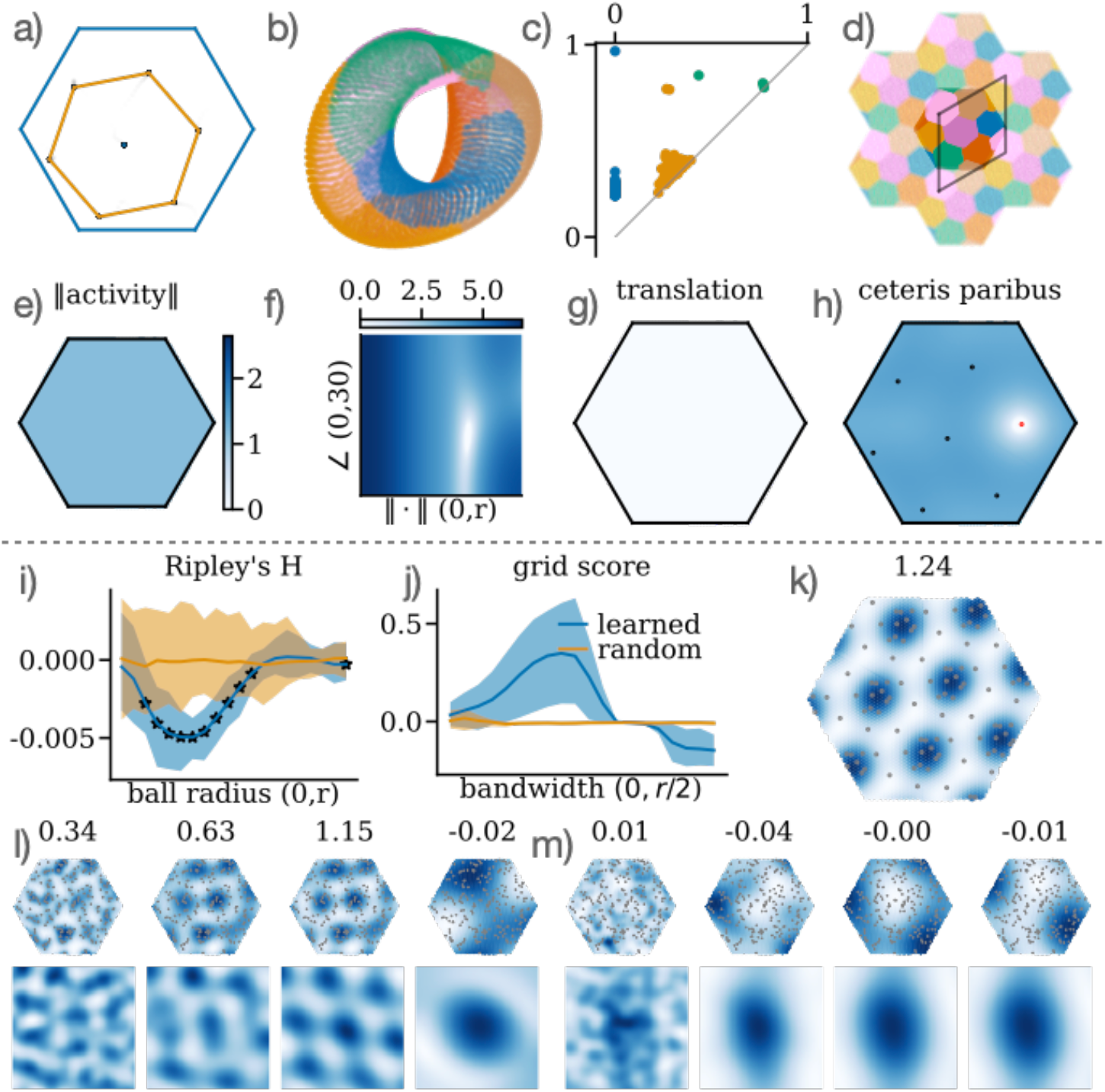
Optimal phases for few (seven, a)-h)) and many (one hundred, i)-m)) grid cells. **a)** Visualization of learned phases for seven grid cells within the unit cell (blue) alongside the inferred hexagonal phase arrangement (orange). **b)** A low-dimensional projection colour-coded by its Voronoi diagram (shown in d)). **c)** A persistence diagram illustrating the activity topology from a). **d)** Voronoi diagram depicting the spatial partitioning of phases across an extended grid of unit cells, with the primitive (rhombus) cell superimposed. **e)** The L2 population activity n orm measured everywhere in the unit cell. The colour bar ranges from 0 to 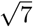, which is the maximum norm a population of 7 cells could generate. **f)** Conformal isometry grid search when varying the angle and magnitude of the hexagonal phase solution in a). Colours are log*C* +1 where *C* is the conformal isometry metric defined in Equation 5. **g)** A common phase translation grid search, maintaining the colour scale from f), to illustrate invariance in phase shifts. **h)** Grid search focusing on varying just one (marked as a red dot) of the phases, using the same colour scale as f).

This expansion leads to two primary consequences: an increase in energy consumption, as measured by the L2 norm of the population activity, and an enhancement in spatial resolution. The increase in energy consumption arises because more cells are actively contributing to encoding locations on the ring, each cell’s activity adding to the overall energy of the system. To quantify the change in resolution, we discretise cell firing rates by assuming an absolute refractory period (e.g., 5 ms) and ignoring subthreshold dynamics, effectively categorising the firing rates into discrete states (e.g., 0, 1, 2, …, 200 Hz). This discretisation transforms the ambient space into distinct voxels, each representing a unique state of the population activity. As the number of cells increases, so does the volume of the state space, the quantity of voxels and the radius of ring. The ring then intersects with a greater number of voxels, thereby including more states, which translates to a higher resolution in the encoded variable. This underscores the trade-off between energy efficiency and the precision of spatial encoding in grid-cell modules.

In addition, a conformal isometry ensures that the encoding is robust to noise. In fact, the error in an encoded variable is locally bounded by a conformal scaling of the error in the neural representation [22]. This can be seen by assuming a noise level *ξ* in the firing rates. We can then differentiate two stimuli, *s*_1_ and *s*_2_ when their distance *d*(*s*_1_, *s*_2_) is at least *ξ*. For a circle, this distance is *d*(*s*_1_, *s*_2_) = *rθ*, where *r* is the circle’s radius and *θ* is the angular separation between *s*_1_ and *s*_2_. Thus, encoding the circle with *N* neurons allows for the differentiation between stimuli separated by a minimum of 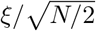 radians.

To make a similar argument for how the surface area of the toroidal manifold of a module of grid cells increases, we analytically derive the conformal scale *σ*_hexagon_. This measure is independent of the phase solution, and given as (see derivation in Conformal scale)

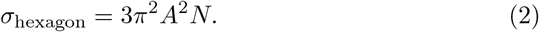

Here, *A* represents the amplitude of the grid pattern, corresponding to the maximum firing rate of each cell, and *N* is the number of cells in the module. Note that this scale constitutes a local minimum, in the sense that it is the scale that minimises (4) irrespective of the phase distribution used.

For comparison, we also derive the conformal scale for a square grid pattern *σ*_square_ and establish that the ratio of the hexagonal to square grid scales is *σ*_hexagon_*/σ*_square_ = 3*/*2. In other words, the hexagonal pattern’s scale is 50% larger than that of a square pattern for the same firing rate amplitude *A* and cell count *N* . Fig. 4c) visualises the conformal scale relative to the cell count and firing rate for the hexagonal pattern, *σ*_hexagon_, with light blue contour lines representing the square grid’s scale, *σ*_square_. That is, resolution and noise-robustness is higher for a module of cells with a hexagonal pattern compared to a module of equally many cells, but with a square pattern.

What implications does a higher conformal scale have in a physical context? First, we find that it appears to maintain constant energy across space, as measured by the L2 norm of the population activity, scaling with the conformal scale (Fig. 4d)). Geometrically, it amounts to inflating the manifold, increasing the surface area of the encoding manifold, as illustrated in Fig. 4g). This corresponds to a finer resolution in the encoded variable (Fig. 4e)) and increases noise-robustness.

Furthermore, our analyses reveal that on average, each phase has to be moved less (from a random uniform initialisation) to achieve a conformal isometry as the number of cells increases (details in Calculation of Learning Trajectory Length. This is illustrated in Fig. 4f)). In this sense, a larger module requires less optimisation to reach a CI. This observation suggests that as the dimensionality of the module increases, the solution space (phase configurations) for conformal isometry becomes denser.

### 2.5 Exploring alternative optimisation objectives

In addition to our primary focus on conformal isometry, we explored two alternative optimisation criteria: (i) minimising linear decoding error and (ii) maximising toroidal persistence. We compared these objectives against a baseline of randomly selected phases as illustrated in Fig. 5. For each criterion, we trained 50 models for 5000 epochs, each with 64 uniformly sampled spatial points and a learning rate of 0.001 and cross-evaluated them against the other objectives.

In the case of linear decoding, the model specifically trained to minimise this error initially outperformed the others. However, as the number of cells increased to seven or more, all models converged to similar performance levels. This convergence suggests that beyond a certain point, merely increasing the number of cells does not improve the performance of a linear decoder. Intriguingly, the linear decoding-optimised model generally performed poorly across other criteria, failing to outperform even the random phase baseline.

The models optimised for toroidal persistence (details in Toroidal loss function) yielded similar CI performance, but achieved a persistent torus for fewer cells compared to a random phase arrangement. On the other hand, models optimised for conformal isometry, not surprisingly, achieved the lowest isometry loss. More surprisingly, however, they also had the most persistent toroidal Figure 4. : **f)** The average geodesic distance between the initial and final positions of phases post-learning, plotted against the number of cells in the module. This metric provides insight into the learning trajectory and its dependence on module size. **g)** A depiction of how increasing the number of cells leads to larger toroidal structures and extended neural trajectories, emphasizing the spatial and topological expansion with larger modules. characteristics for seven cells and upwards, although the gap to random phases and homology-optimised phases decreases with more cells.

## 3 Discussion

In this work, we explore the capacity of idealised grid cells with freely varying phases to form a conformal isometry (CI) of two-dimensional flat space. We discover that as few as seven cells are sufficient to achieve such a spatial encoding. This encoding allows the module to uniquely represent its unit cell, form a torus, and form a conformally isometric map to flat two-dimensional space. Remarkably, the phases of these seven cells arrange into a hexagonal lattice, a pattern that persists and extends as the number of cells in the module increases. This arrangement enhances spatial resolution and noise robustness throughout the unit cell, albeit with increased energy consumption, highlighting a trade-off inherent in the system’s precision and energy cost.

**Figure 4.**
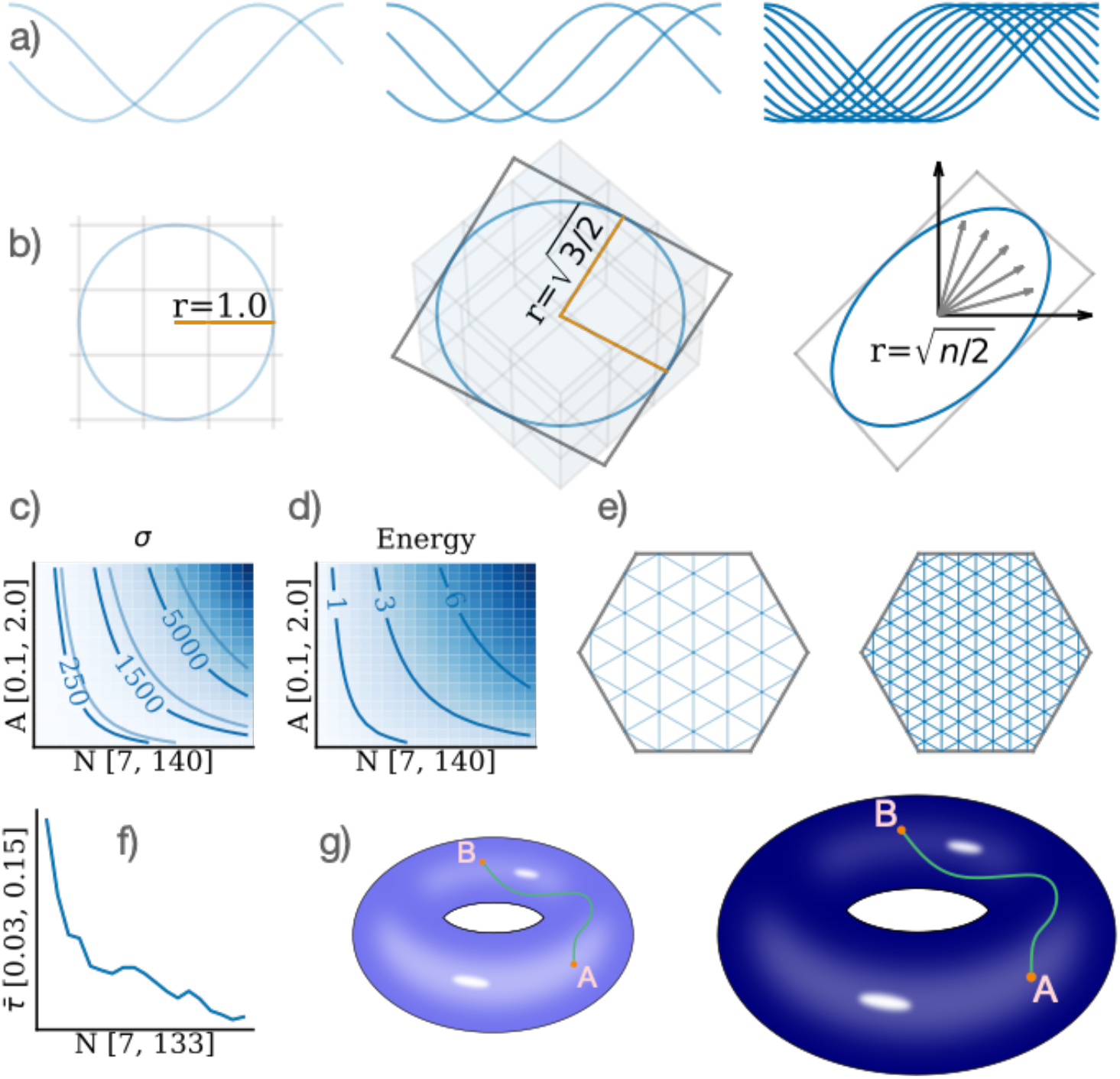
Impact of the Number of Cells on Spatial Encoding. **a)** Tuning curves for 2, 3, and 10 neurons demonstrating one-dimensional periodic encoding. **b)** Population activity forming a ring with expanding radius as the number of cells increases. **c)** The conformal scale (*σ*_hexagon_) plotted against the firing rate (*A*) and number of cells (*N*), illustrating the relationship between module size and spatial encoding scale. Light blue contour lines represent the scale for a square grid pattern (*σ*_square_), providing a comparative reference. **d)** The energy consumption of the module, measured as the L2-norm, is shown relative to the firing rate and number of cells. This graph illustrates how energy requirements scale with module size and activity. **e)** A schematic of the unit cell with overlaid meshes of different granularities to depict spatial resolution. The left side, with a coarser mesh, represents a module with fewer cells, while the right side, with a finer mesh, represents a module with more cells, highlighting the resolution enhancement with increased cell count.

**Figure 5.**
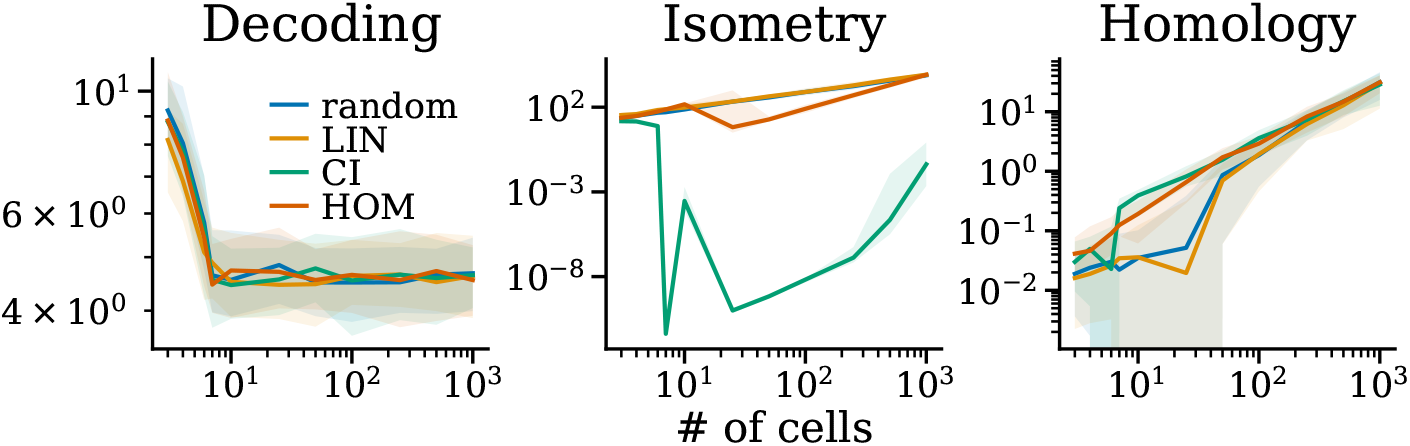
Model comparison: Three models, linear decoding (LIN), conformal isometry (CI) and homology (HOM), along with random phases, evaluated on each other’s metrics.

The demonstration that a module of idealised grid cells can indeed form a CI supports the notion that this property is plausible and potentially fundamental for generating a hexagonal pattern, as suggested in recent studies [21]– [24]. This naturally aligns with the proposition that grid cells provide a neural metric for space [1], [15], [25]–[30]. In fact, by being conformally isometric to flat space, the module locally inherits all the properties of the Euclidean metric. Global distances can subsequently be computed by path-integrating neural trajectories. Furthermore, the observation that a CI solution maintains constant energy throughout space underscores the potential role of energy constraints. In particular, we propose a constant population activity norm across space as an energy constraint for a module of grid cells, similar to the constant unit normalisation in [21], [23]. Together, our results contribute to the understanding of constraints in recent normative models employing recurrent neural networks trained for path integration to learn grid-like solutions, explain how grid cells can form a metric for space.

Interestingly, while a random phase arrangement typically requires around 20 cells to form a torus consistently [38], a seven-cell module optimised for conformal isometry also achieves this, as evidenced in Fig. 3b) and c)). This places the minimal set of grid cells needed for the emergence of a torus (6 cells) in a curious position between the minimal set of grid cells needed to represent the unit cell (3 cells), and representing space equally everywhere (7 cells), leading to an open question about the intrinsic value of having a toroidal representation.

In our investigation of alternative objectives, we explored the concept of encoding spatial positions within a toroidal neural representation and evaluated the possibility of linearly decoding this information from grid cell modules. Our research indicates that a minimum ensemble of six grid cells is necessary for accurate toroidal encoding, highlighting the intrinsic complexity of spatial representations in neural systems. Furthermore, we found that the linear decoding of Cartesian coordinates from grid cell activity faces significant limitations, suggesting a potential mismatch between traditional linear decoding models and the actual grid cell encoding mechanisms. This insight calls for a reevaluation of existing normative models, advocating for a more nuanced understanding of how grid cells contribute to spatial cognition and navigation.

Despite allowing for theoretical insights, the simplicity of our model does not account for several experimentally observed features of the grid cell system, restricting its explanatory scope. For example, it would be interesting to study whether a CI in environments of different geometries or with salient landmarks could explain distortions in the grid pattern [28], [47]–[49], e.g., by locally adjusting the CI scale. Furthermore, while grid cells may be involved in performing other tasks, Xu *et al*. show that CI also supports path integration [21] and the model could be readily extended to map different spaces [50], [51]. Including interactions between multiple grid cell modules or decoders of greater expressivity could also improve the decoding of Cartesian positions from a grid cell module.

The simplicity of our model, however, enables testable predictions about how a single parameter (grid phases) relates to the presence of a CI and opens up the path for experimental validation. For instance, large-scale recordings within a module of grid cells could be used to test whether grid cells set up a conformal isometric map of space and whether their phases arrange in a regular and hexagonal manner.

## 4 Methods

### Grid cell model

We model grid cells following the approach of Solstad *et al*. [31], representing grid cells as a superposition of three plane waves with wave vectors *k*_*j*_, each offset by 60 degrees. The model can be expressed as:

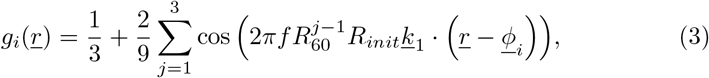

where *i* indexes grid cells, *R*_*init*_, *R*_θ_ ∈ *SO*(2) are rotation matrices determining the grid pattern orientation offset and the relative orientation between wave vectors, respectively. *f* represents the spatial frequency while *<u>r</u>* ∈ℝ^2^ and ϕ_*i*_ ∈ ℝ^2^denote the spatial and phase coordinates, respectively. We set *f* = 1, and by applying the rotation matrix *R*_60_ we get the three wave vectors, 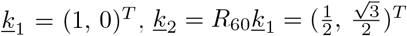, and 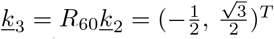. The phases are specified in the relevant parts of the text (typically initially random uniform).

### Conformal isometry loss function

To achieve a conformal isometry, we introduce a loss function ensuring that distances in neighbouring points in 2D space are proportional to distances in the grid code. The loss function is defined as:

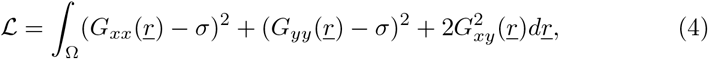

where, *σ* is the isometry scale, and *G*_*xx*_, *G*_*yy*_, and *G*_*xy*_ are components of the metric tensor *G* = *J*^*T*^ *J ∈* ℝ^2*×*2^. Here, 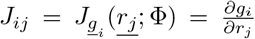 denotes the Jacobian matrix derived from differentiating Equation (3) with respect to the position *r* for a fixed set of *N* phases Φ∈ℝ^*N×*2^. The notation emphasises the Jacobian’s dependency on the phase configuration. The integral over the domain *Ω*, which represents the unit cell of the grid pattern, is approximated using Monte Carlo integration by uniformly sampling points within the domain.

### Conformal Isometry Evaluation

To evaluate the degree of conformal isometry achieved by the grid module without explicitly setting the isometry scale we focus instead on the variance and expected values of the metric tensor components over a set of spatial samples. We define the conformal isometry score (CIS) as follows:

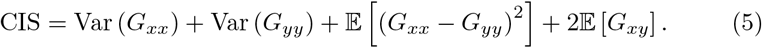

This metric encapsulates the deviation from an ideal conformal isometry, where the variance of *G*_*xx*_ and *G*_*yy*_ would be minimised, and the expected values of the off-diagonal component *G*_*xy*_ and the difference between *G*_*xx*_ and *G*_*yy*_ would approach zero. A lower CIS value indicates a closer approximation to a true conformal isometry, reflecting a more uniform and isotropic representation of space by the grid module. This provides another way to evaluate the quality of the spatial encoding to form a conformal isometry, without knowing the conformal scale.

### Toroidal loss function

To define a loss function which maximises the topological signatures of a torus, which are two one-dimensional holes and one two-dimensional hole, we followed the approach outlined in Gabrielsson *et al*. [52] and used the `torch_topological` python package [53]. From a grid cell module with randomly initialised phases, we compute the pairwise distances between grid activities and use them to construct a Vietoris-Rips filtration as a function of the scale parameter *ϵ*. The persistence of a topological feature *ω* is defined as the point at which the feature disappears (the death) *d*(*ω*) minus the point at which it first appears (the birth) *b*(*ω*),

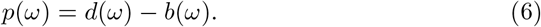

If we then sort all features of dimension *d* in ascending order according to their persistence *{p*_*d*_(*ω*_1_), …, *p*_*d*_(*ω*_*N*_)*}*, we can define our loss function as

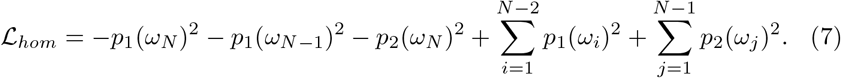

As one can see, this loss is minimised when the two largest one-dimensional and the largest two-dimensional features are maximised, and all other features disappear.

### Linear Euclidean decoding

To find phases that are optimal for linear decoding we jointly optimised both the phases Φ and a linear readout matrix *W* of shape *N ×*2 for reconstructing a target Cartesian representation of positions *r ∈ Ω*. The weights *W* were initialised as 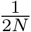and the phases were initialised randomly uniformly in the unit cell *Ω*. We start by creating a *T ×*2 dimensional (response) matrix *P* of positions sampled uniformly from the unit cell Ω. Subsequently, we construct a corresponding *N×T* (design) matrix *X* of grid cell activities *X*_*ij*_.= *g*_*j*_ (*r*_*i*_) The weight matrix *W* and the phases Φ were optimised to minimise the loss

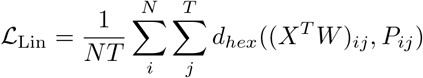

where *d*_*hex*_ is as described in Shortest distance on hexagonal unit cell.

### Training details

Uniform spatial samples <u>*r*</u> in the unit cell Ω are generated via rejection sampling using a minimum enclosing square of the unit cell as the proposal distribution. We use the Adam optimiser [54] with default PyTorch [55] parameters and mini-batches of size 256 of spatial samples *r*, except where specified otherwise. Training length varies between 500 and 10000 training steps for different experiments and is specified throughout the text. We choose training lengths to achieve a small loss, and we observe that models with more cells typically converge faster and, thus, require fewer training steps.

### Low-dimensional projection

We use UMAP [56] to project the activity of multiple grid cells down to three dimensions (*n_components* = 3). We modify default parameters for Fig. 1e) by setting *n_ neighbours* = 25. For Fig. 2c) we set *n_ neighbours* = 40 and *min_ dist* = 0.2. For Fig. 3b) we use *n_neighbours* = 1000.

### Persistent cohomology

All persistence diagrams are computed using the Ripser package [57] with default parameters apart from setting *maxdim* = 2 and *n _perm* = 150.

### Ripley’s H-function

We use Ripley’s H function [44]–[46] to analyze phase dispersion or clustering over various scales within a hexagonal domain. It is computed by first wrapping all phases into their unit cell. Then, the phases are duplicated to their six immediately neighbouring hexagons to handle edge effects (see Fig. 3d) for an example of the extended tiling). Subsequently, balls with a given radius *ϵ* are centred at each (non-duplicated) phase ϕ _*i*_ . The core of this analysis involves counting the number of phase points encompassed within each of these balls and normalizing the count. This procedure is mathematically formulated as Ripley’s K-function, represented by:

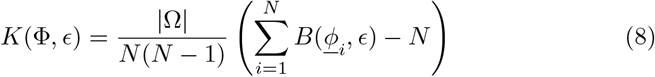

where *B*(.) signifies the ball counting function (which counts the number of phases within a ball), *N* is the count of original (non-duplicated) phase points, and |Ω | denotes the area of the unit cell.

Ripley’s K function can be extended to have zero mean, yielding Ripley’s H-function, expressed as

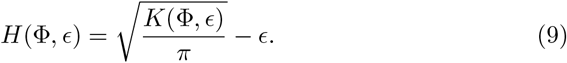

### Permutation test

In the permutation test, we evaluate whether the response values of two groups, *A* and *B* significantly differ. The test aims to determine the similarity or dissimilarity between these groups using a specific statistic. In our case, we use the sample mean difference, denoted 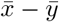, as the test statistic.

The procedure begins by calculating this statistic between groups *A* and *B*. We then merge the samples from both groups and randomly draw two new samples, *A*^*′*^ and *B*^*′*^, from this combined pool. The test statistic is recalculated for these new samples. This process of permutation and recalculation is repeated 200 times (*n*_*perms*_ = 200), and the resulting statistics are ordered to determine the two-sided percentile (p-value) of the original, non-permuted statistic. The null hypothesis, which posits “no difference”between the groups, is rejected if the p-value is less than the significance level of α = 0.01.

Group *A* consists of Ripley’s H scores of 50 CI-optimised models, each with 100 cells. Each model is trained from a random uniform phase initialisation for 500 mini-batches of size 256. The loss for these models and parameters typically decreases from an initial value of around 10^*−*1^ to between 10^*−*5^ and 10^*−*8^. Group *B* comprises 50 models with random uniformly arranged phases within the unit cell, with its sample size matched to that of group *A* (|*B*| = |*A*| = 100).

The permutation test thus provides a way to evaluate whether the optimised phases in group *A* differ significantly from the random uniform distribution in group *B*, based on Ripley’s H statistic.

### Phase KDE

To analyse the distribution of phase points, we apply a kernel density estimate (KDE) using a Gaussian kernel. Initially, we wrap the phases into their unit cell. To capture the long-range periodic effects, we then duplicate the phases to a double-tiling of the unit cell. For the KDE calculation, we utilise the Scipy implementation scipy.stats.gaussian kde [58]. The bandwidth parameter bw method is detailed in the description of each analysis.

### Grid score

To assess the hexagonal symmetry in the phase distributions, we employ a modified grid score computation. The process begins with the calculation of the Gaussian phase KDE, as detailed in Phase KDE. This KDE is then evaluated over two square mesh grids: a smaller grid covering the interval [*−r, r*]^2^ in ℝ^2^ with a resolution of 64 *×*64 pixels, and a larger grid spanning [*−*3*r/*2, 3*r/*2]^2^ with 127*×*127 pixels. These grids, or ratemaps, are spatially correlated using the Pearson correlation coefficient in ‘valid’ mode, producing an autocorrelogram of 64 × 64 pixels. This autocorrelogram corresponds to the minimal enclosing square of the unit cell, maintaining the periodicity of the pattern.

Next, we construct a binary annulus mask matching the autocorrelogram’s dimensions. The outer circle of the annulus is defined by the unit cell’s radius, while the inner circle’s width is set equal to the KDE kernel bandwidth, as specified in the text. This masked autocorrelogram is then correlated with rotated versions of itself at angles of 30, 60, 90, 120, 150, and 180 degrees, again using the Pearson correlation coefficient.

The final grid score is derived by calculating the mean difference between the correlation coefficients at angles of 60, 120, 180 degrees and those at 30, 90 and 150 degrees. This method effectively quantifies the degree of hexagonal symmetry present in the phase distributions.

### Hexagonal unit cell mesh

To create a hexagonal mesh, we start by generating a square mesh of spatial points *r* within the interval [0, 3*R/*2]^2^, where *R* is the radius of the unit cell. This mesh is initially conceptualised in a 60-degree rhombus basis relative to the standard basis. We invert this representation to the standard basis with the inverse rhombus transform *T*^*−*1^. The basis change matrix is, thus, given by *T* = (*<u>e</u>*_1_,*R*_60_*<u>e</u>*_1_)^*T*^ where *<u>e</u>*_1_ = (1, 0)^*T*^ and *R*_60_ a 60-degree (counter-clock wise) rotation matrix. Finally, we wrap the corresponding rhombic coordinates to the hexagonal unit cell.

### Shortest distance on hexagonal unit cell

To determine the shortest distance between two points, *<u>r</u>*_1_ and *<u>r</u>*_2_, within a hexagonal unit cell with periodic boundaries, we select one point *<u>r</u>*_2_ and replicate it in all nearest neighbour unit cells. The shortest distance respecting periodic boundary conditions is acquired by selecting the shortest out of all distances between *<u>r</u>*_1_ and *<u>r</u>*_2_ including the six replicas.

### Voronoi Diagram Construction

We construct a Voronoi diagram, categorising each spatial point of a Hexagonal unit cell mesh. Each point is categorised to belong to the phase that it has the Shortest distance on hexagonal unit cell to.

### Calculation of Learning Trajectory Length

To quantify how the phases evolve during training, we calculate the learning trajectory length. This metric represents the average distance travelled by the phases from their initial positions <u>ϕ</u> ^0^ to their final positions at the end of training <u>ϕ</u>^T^. We define the learning trajectory length 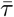 as:

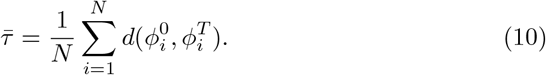

*·*Here, *N* is the number of phases, and *d*(.) represents the distance function as defined in Shortest distance on hexagonal unit cell. By averaging these distances across all phases,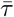effectively captures the overall extent of movement or adjustment the phases undergo during the learning process.

### Author Contributions

VSS Developed code, performed experiments and analysis and wrote the paper.

KB Developed code, performed experiments and analysis and wrote the paper.

MBP Developed code, performed experiments and analysis and wrote the paper.

EH Developed code, performed experiments and analysis and wrote the paper.

KH Developed code, performed experiments and analysis and wrote the paper.

AMS Supervised the process and wrote the paper. MF Wrote the paper. MEL Supervised the process and wrote the paper.

## Acknowledgements

This research was funded by the Research Council of Norway Grant 300504, the University of Oslo and Simula Research Laboratory.

## Data and code availability

All original code has been deposited at https://github.com/bioAI-Oslo/CI-grid-cells and is publicly available as of the date of publication. Any additional information required to reanalyse the data reported in this paper is available from the lead contact upon request.

## Appendix

### A Loss history

In Fig. A1, we track the conformal isometry loss (from Equation 4) over 5000 training iterations for cell counts ranging from 1 to 14. Notably, there is a stark reduction in loss, approximately ten orders of magnitude, when utilizing seven cells compared to fewer. This contrast suggests that a module of seven grid cells is a threshold for forming a conformal isometry, where fewer cells can not.

Further observations reveal that cell counts from 8 to 14 generally achieve similar losses, albeit with variations in the number of training iterations required. However, modules with 10 and 12 cells exhibit peculiar behaviours; particularly, modules with 10 cells seem unable to reach the lower loss magnitudes indicative of an accurate conformal isometry. This may hint at an inherent limitation or suboptimal configuration within the 10-cell arrangement for encoding space as a conformal isometry.

### Seven colour map of hexagonal torus

Figure. A2 shows an example of the seven-colour theorem.

### Toroidal topology for CI optimised models

In order to answer the question of how many cells are needed for a group of grid cells to generate a torus, we trained CI models with between one and fourteen cells. In terms of (co)homology groups computed through persistent (co)homology, a torus is characterised by having two large (persistent) one-dimensional features and one two-dimensional feature.

As one can see from Fig. A3, which shows the sizes of these features as a function of the number of cells, the two-dimensional feature appears at three cells. At five cells there are both a two-dimensional and a one-dimensional feature, but the second two-dimensional feature is yet to appear. At six cells, there is a qualitative jump in the second one-dimensional feature, at this point the manifold has all the features necessary to characterise a torus. We therefore claim that six cells is an upper bound on the minimum amount of cells necessary for the emergence of a torus. This observation is in line with the empirical fact that one needs at least 6 PCA components to account for most of the variance in grid-cell data. A theoretical explanation for why this might happen can be found in the supplementary information of Gardner *et al*. [37].

### Ripley’s H sanity checks

To verify the results of the Ripley’s H function presented in Fig. 3i), we conduct a series of sanity checks. We start with a model of seven cells optimised for conformal isometry over 5000 iterations. From this optimised model, we infer the learned hexagonal solution and replicate it 15 times, introducing varying degrees of noise (common phase shift) to each copy.

In the left panel of Fig. A4, we introduce normal noise with zero mean and three distinct spreads, *σ*, to the hexagonal solution. For a minimal spread of *σ* = 0.01, the Ripley’s H function exhibits a highly deterministic and pronounced zig-zag pattern. This observation is facilitated by 40 permutations of noise for each spread, allowing us to include *±* standard deviation error bars. The zigzag pattern arises because, with minimal noise, so that even balls of small radius encompass most points, resulting in a higher-than-expected number of points within these balls. As the radius increases without including additional points, the function dips, reflecting a decrease in the relative expectation of points per ball. However, once the radius extends slightly beyond 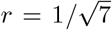, the characteristic radius of the hexagonal Voronoi cells (refer to Fig. 3d)), it suddenly includes phases from adjacent clusters, causing a sharp uptick in the number of points within the balls.

Conversely, as the noise spread increases to *σ* = 0.05 and 0.15, Ripley’s H function transitions to a smoother and flatter profile. This change is logical, as a larger spread of noise means that balls must be larger to encompass the more dispersed points, leading to a more gradual change between the expected and actual number of points within the balls. This smoothing effect reflects the increased randomness and dispersion of points, diluting the pronounced zig-zag pattern observed at lower noise levels.

The right panel of Fig. A4 contrasts a baseline of random uniform sampling within the unit cell (green curve) against the structured arrangement of the hexagonal solution (orange curve). Here, 15 copies of the 7-cell hexagonal solution are subjected to noise, which, unlike the left panel, is uniformly distributed across the hexagonal cell, leading to completely random centres for the copied solutions. Despite this randomness, the inherent structure of the hexagonal solution imparts a distinct spatial statistic, as evidenced by Ripley’s H function. This distinction is particularly noticeable as a dip in the graph be low the excepted number of points for balls with a radius smaller than 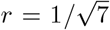. Subsequently, we can see a sharp transition to a higher-than-expected number of points as the radius of the balls includes neighbouring hexagonal points. This characteristic is markedly evident in the hexagonal arrangement, where the surrounding points abruptly contribute to the count, creating a clear demarcation in the Ripley’s H function. These transitions can be somewhat smoothed out for non-perfect hexagonal copies, leading to a less pronounced but still distinguishable pattern compared to a completely random phase distribution, as also observed in Fig. 3i). This behaviour underscores the unique spatial signature of the hexagonal arrangement, even amidst increased randomness.

### Independence of CI solutions

Our analysis reveals that if a grid cell population 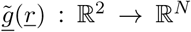forms a solution to the conformal isometry loss function (Equation 4) with scale 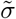, then an extended population that includes a phase-shifted copy of this population also forms a solution, with the scale doubled to 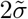.

To demonstrate this, consider the extended population vector defined as:

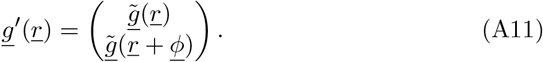

The Jacobian of this extended population can be expressed as a block matrix:

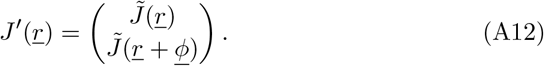

This formulation allows us to derive the metric tensor of the extended population:

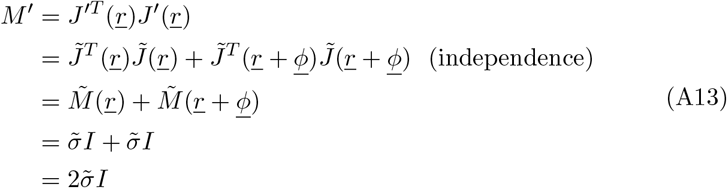

where we in the second step have used that a matrix *X ϵ ℝ*^*m×n*^ multiplied with itself *X*^*T*^ *X* can be written as (*X*_1:*m/*2_)^*T*^ *X*_1:*m/*2_ +(*X*_*m/*2:*m*_)^*T*^ *X*_*m/*2:*m*_.

In the fourth step, we are using that a module of grid cells that constitute a CI is independent (still a solution to CI) to a common phase shift. This can be seen by inducing a common phase shift *<u>ϕ</u>* in each grid cell and rewriting it as a spatial displacement of the grid module *g*(*r* + *<u>ϕ</u>*).. By virtue of CI, the metric tensor of *<u>g</u>* is constant for all *<u>r</u>*, vec *<u>r</u>* + *<u>ϕ</u>*. Hence, a common phase shift to a CI solution is also a CI solution. An example of this is shown computationally in Fig. 3g).

### Conformal scale

The conformal isometry loss is given by (4). We want an extremum, which is given by 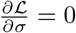, which yields (assuming the integrand is well behaved)

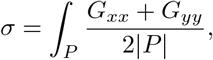

where |*P*| is the volume of the region of integration, which is the parallelogram spanning the pattern unit cell. For the second derivative, we have that

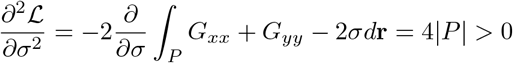

Thus the value of *σ* is a (local) minimum for a nonzero region of space.

Note that the parallelogram region spans all six hextants of the hexagonal unit cell of a grid pattern of radius *<u>r</u>*. However, we will rather perform integration in the unit square *U* . The transformation between the two is given by

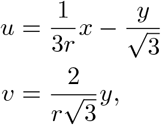

where (*x, y*) are parallelogram coordinates, and (*u, v*) square coordinates. Then, when *U* is the unit square,

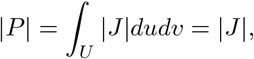

with *J* being the jacobian of the transformation. For the scale *σ*, we have that

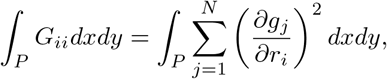

but we integrate over the full domain of each *g*_*j*_, which are all of the same functional form, so every contribution is equal. Thus

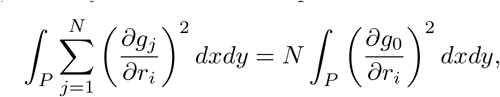

where *g*_0_ is a grid function centred at the origin (zero phase shift). Note that this step corresponds to a coordinate shift removing the phase in each term.

### Hexagonal grid Functions

A hexagonal grid function can be constructed using three plane waves, with wave vectors offset by 60 degrees. In other words, we may write a zero phase shift grid function as

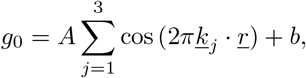

and

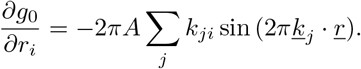

We set the wave vectors to be

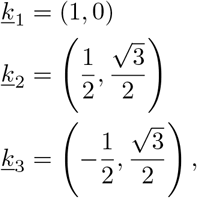

So

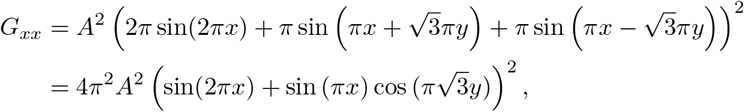

where we have used the identity sin(*u*+*v*)+sin(*u−v*) = 2 sin(*u*) cos(*v*). Likewise

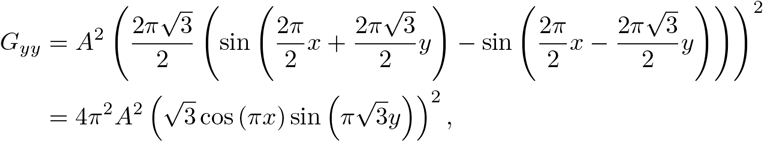

where we have used that sin(*u* +*v*) *−* sin(*u − v*) = 2 cos(*u*) sin(*v*). We thus need to integrate

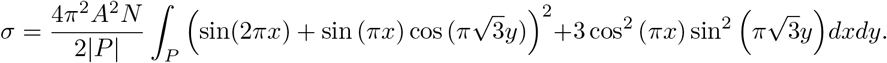

We then perform the coordinate transformation, and insert the Jacobian determinant (which cancels |*P* |), and find that the integral may be written

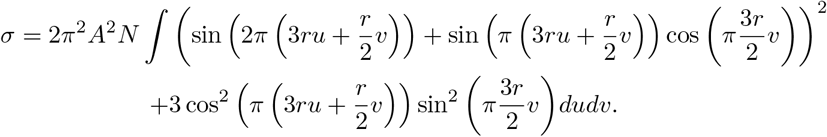

which, using an integral solver, yields

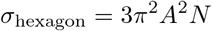

### Square grid functions

The derivation is the same for a square grid pattern, only simpler. A square grid can be constructed using two plane waves, whose wave vectors are offset by 90 degrees. Thus

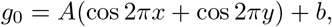

and

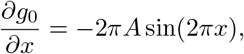

and

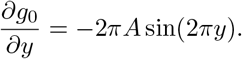

Then

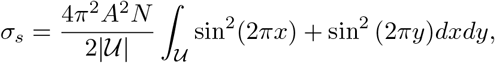

Where |*𝒰*| = 1 is the area of the square unit cell. Then

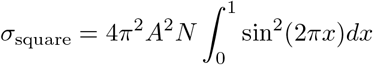

which is just

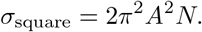

### Hexagonal vs. Square scale

We have that

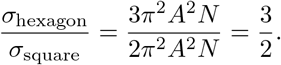

### Proof of Injection

We consider

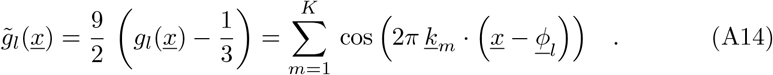

similar to Equation (3) which expresses the grid field of a single grid cell characterised by the *K* wave vectors {*k*_*m*_}. We will take a look at some examples with concrete realisations of *k*_*m*_ later. Each grid cell *l* is characterised by a phase vector *<u>ϕ</u>*_*l*_ . In accordance with behavioural experiments, we assume a 2-dimensional world, and hence *<u>x</u>* ∈ ℝ^2^. We collate the population response of *N* grid cells in one *N* -dimensional vector *H* ∈ [-1, 1)^*N*^ ⊂ ℝ^*N*^

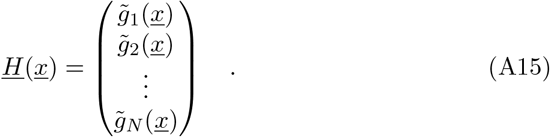

#### Theorem A.1

(Unique Grid Cell Code). *A bijective grid cell code of K wave vectors {k*_*m*_*} requires at least N* = *K grid cells with mutually different phases {<u>ϕ</u>*_*l*_.*}*

*Proof*. First, we introduce the complex formulation of the problem by using Euler’s formula and taking the real part of the expression to ensure *H* remains real-valued

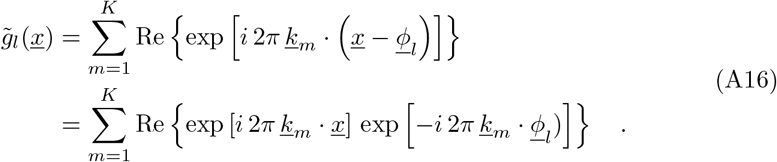

Since taking the real part is a linear operation which preserves linear structures,

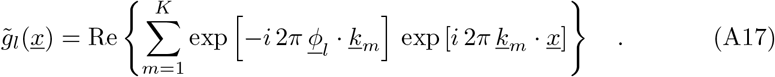

For convenience, 1we introduce matrix notation. Let *A ∈* ℂ^*N×K*^ with *A*_*lm*_ =exp[−*i* 2π *<u>ϕ</u>*_l_·*km*] and υ ∈ ℂ^*K*^with υ*m*.(*x*)= exp [*i* 2π*k*_*m*_ ·*x*]. Then, we notice that Equation A17 formally expresses a matrix multiplication such that

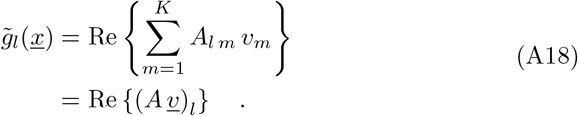

Consequently, the population grid field becomes

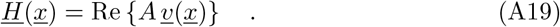

In order for *<u>H</u>* (*<u>x</u>*) to be locally bijective (one-to-one and onto), a neighbourhood *U ⊂* ℝ^2^ needs to exist such that for every 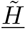 in a neighbourhood *V ⊂* [*−*1, 1)^*N*^ exists exactly one *x* ∈ *U* such that

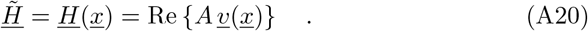

Assuming such neighbourhoods exists, Equation (A14) constructs a respective 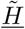 for each *x* ∈ *U* . In order to show that the grid field is one-to-one, we need to provide a way of constructing a corresponding *x* for each 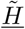. To do so, weneed to resolve Equation (A20) for υ. Equation (A20) can be resolved iff a pseudo-inverse *A*^+^ ∈ ℂ^*K×N*^ for *A* exists such that *A*^+^ *A* = 𝕀. That is the case iff *A* has full rank: rank(*A*) = min(*K, N*); and rank(*A*) = *K*. Consequently, *A*^+^exists iff

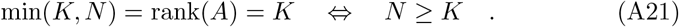

Consequently, the grid cell code can only be bijective if we have a response from at least *K* different grid cells. Because the entries in *A* are determined by the dot products *<u>ϕ</u>*_l_ . *k*_m_, full rank requires at least *K* mutually different phase vectors *<u>ϕ</u>*_l_. Since 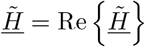 and taking the real part is linear,

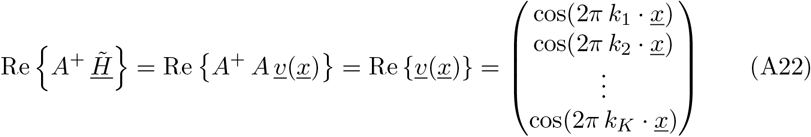

For every *Q*_*m*_ = [2*π k, π* +2*π k*) or *Q*_*m*_ = [*−π* +2*π k*, 2*π k*) with arbitrary *k ∈* ℤ, the cosine function is bijective on *Q*_*m*_.2π *k*_*m*_ . *x* ∈*Q*_*m*_ iff

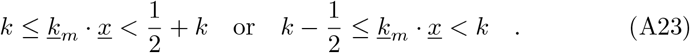

For both cases exist open neighbourhoods *U*_*m*_ *⊂* ℝ^2^ such that the respective criterion is satisfied ∀ *x*.∈ *U*_*m*_ Consequently, the grid cell code is bijective in 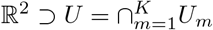.

### Trainable Conformal Scale

To verify the scaling law in (2), we trained multiple models with varying numbers of grid cells within a module to minimise the CI loss in (4), treating the conformal scale *σ* as a trainable parameter alongside the phases. Each model was trained for 10000 training steps using gradient descent and the Adam optimiser with a batch size of 64 spatial samples drawn uniformly within the unit cell of the grid pattern. For training, we used a learning rate of 0.01 and otherwise standard optimiser parameters.

Learned scale parameters are shown in Fig. A5a), alongside the corresponding analytic scale. Notably, learned scale parameters match their analytic counterparts across module sizes. This suggests that the analytical conformal scale derived in Conformal scale indeed corresponds to a local minimum of the conformal isometry loss.

Note that training was more unstable when the conformal scale was left trainable. This motivated us to fix the scale parameter according to the scaling law in (2) when optimising other models. However, the trainable-scale models also achieved highly uniform CI-like solutions. For reference, Fig. A5b) shows the metric components for a trained model with 100 grid cells. The diagonal metric components are both spatially uniform (up to two decimal places) and almost equal, while off-diagonal components are near-zero everywhere.

**Figure A1:**
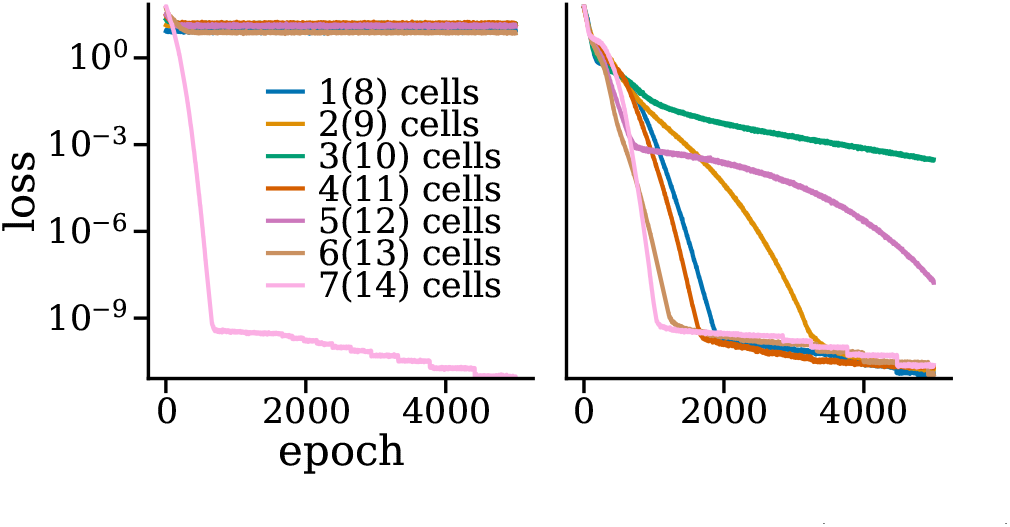
Conformal isometry loss history for 1-7 cells (left figure) and 8-14 cells (right figure).

**Figure A2:**
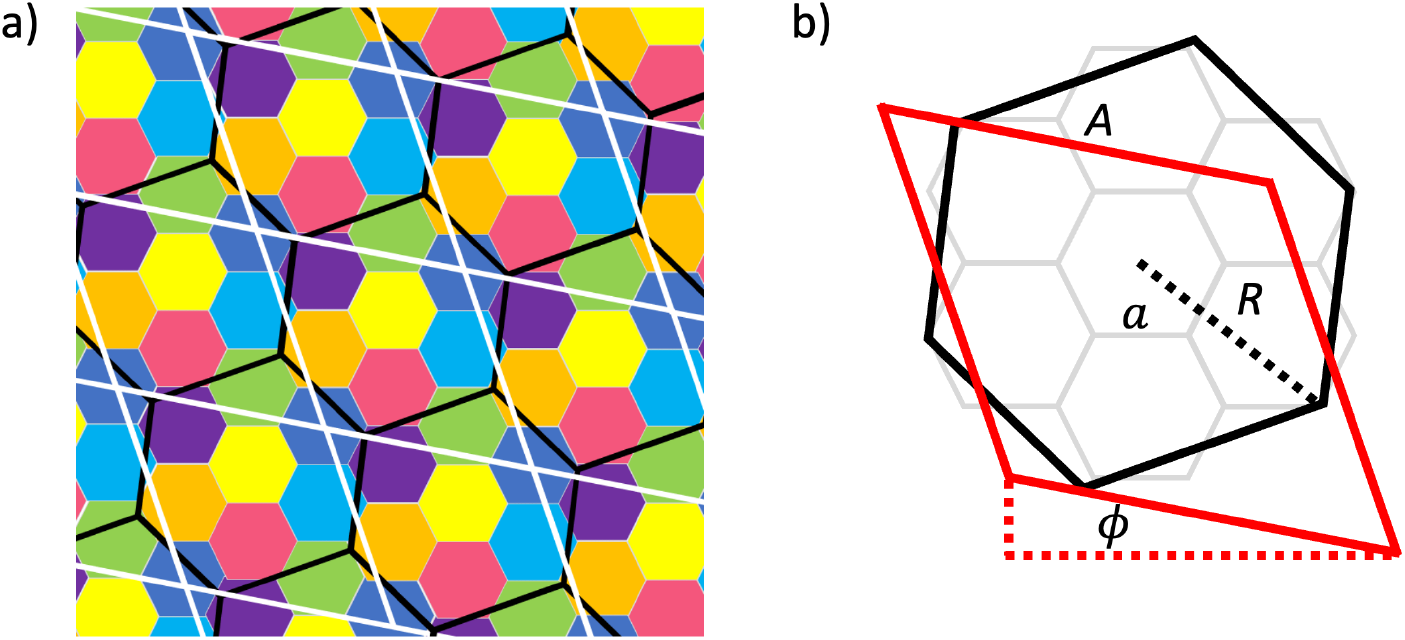
Visualisation of the 7-color map of a flat hexagonal torus. **a, b)** 7 hexagons (coloured/grey), with side lengths *a*, optimally and recursively arranged to tile a plane, are contained in repeating, larger hexagons (black) of radius 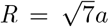 or rhombi (white/red) of side lengths 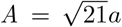, tilted 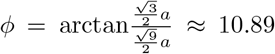 degrees compared to the orientation of the smaller hexagons.

**Figure A3:**
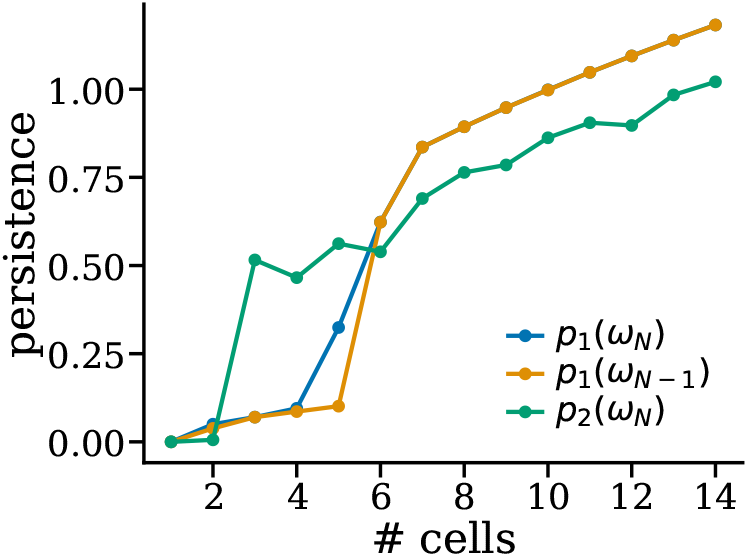
Persistence of the largest one dimensional feature (blue), the second largest one dimensional feature (orange) and the largest two dimensional feature (green) in a CI models with different numbers of cells.

**Figure A4:**
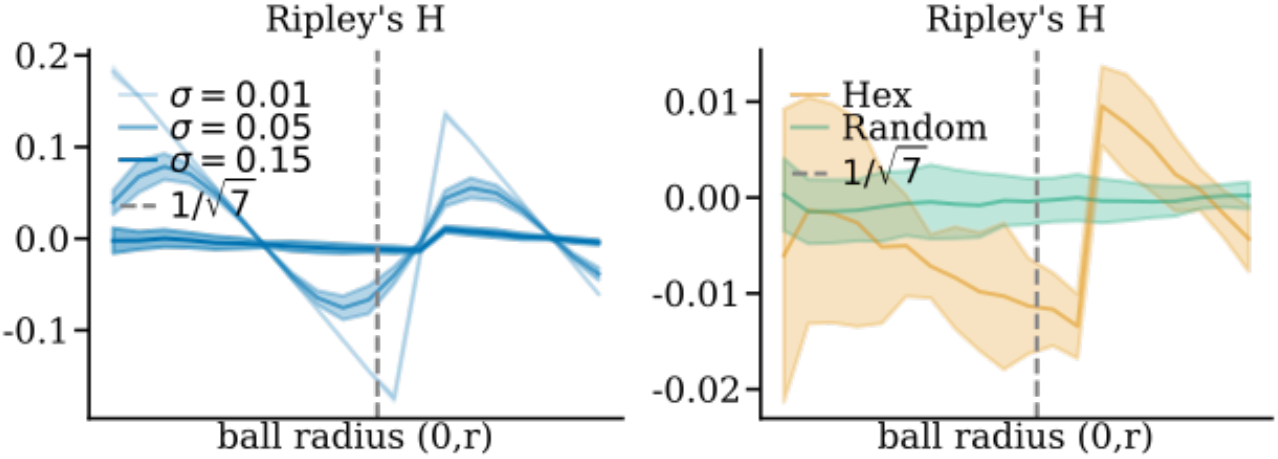
Ripley’s H function for 15 copies of a 7-cell hexagonal solution with varying common noise shifts.

**Figure A5:**
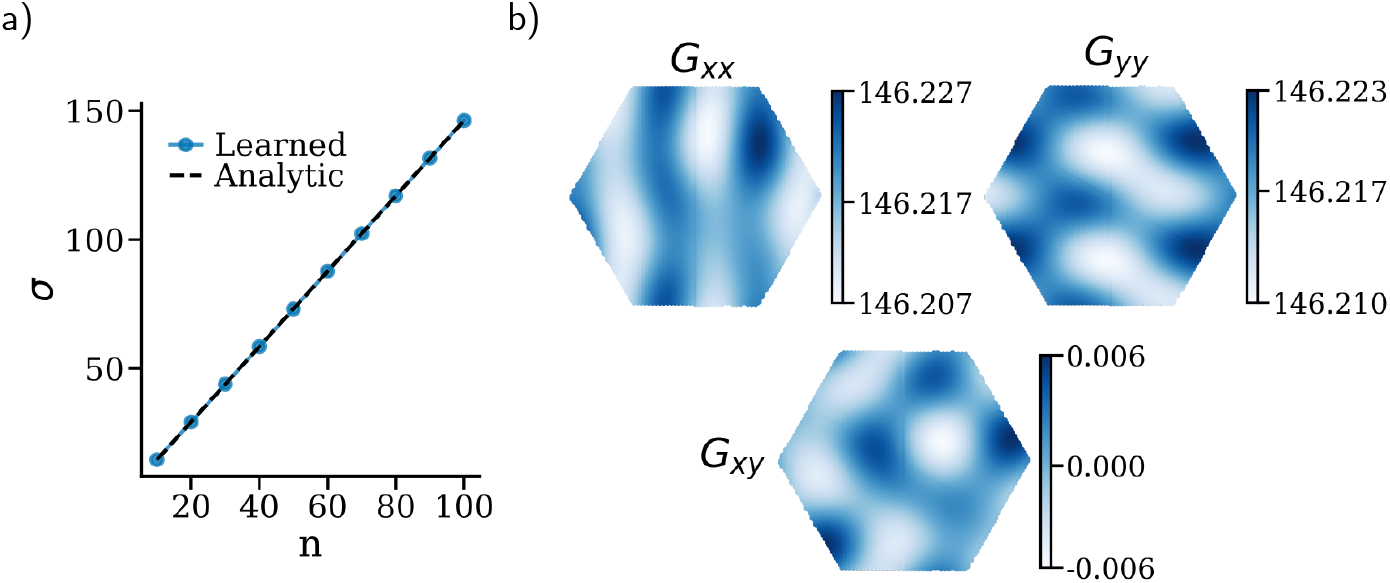
Verifying the conformal scaling law. a) Learned conformal scale (*σ*) as a function of the number of grid cells (*n*) in the module. Also indicated is the theoretically predicted scale (dashed line). b) Metric components for a module of *n* = 100 grid cells with trainable phase distribution and scale.

## References

[1] T. Hafting, M. Fyhn, S. Molden, et al., “Microstructure of a spatial map in the entorhinal cortex,” Nature, vol. 436, no. 7052, pp. 801–806, Aug. 2005. doi: 10.1038/nature03721.

[2] M. Fyhn, S. Molden, M. P. Witter, et al., “Spatial representation in the entorhinal cortex,” Science (New York, N.Y.), vol. 305, no. 5688, pp. 1258–1264, 2004. doi: 10.1126/science.1099901. eprint: https://www.science.org/doi/pdf/10.1126/science.1099901.

[3] Y. Burak and I. R. Fiete, “Accurate Path Integration in Continuous Attractor Network Models of Grid Cells,” PLoS Computational Biology, vol. 5, no. 2, O. Sporns, Ed., e1000291, Feb. 2009. doi: 10.1371/journal.pcbi.1000291.

[4] B. L. McNaughton, F. P. Battaglia, O. Jensen, et al., “Path integration and the neural basis of the’cognitive map’,” Nature Reviews Neuroscience, vol. 7, no. 8, pp. 663–678, 2006.

[5] M. C. Fuhs and D. S. Touretzky, “A spin glass model of path integration in rat medial entorhinal cortex,” Journal of Neuroscience, vol. 26, no. 16, pp. 4266–4276, 2006. doi: 10.1523/JNEUROSCI.4353-05.2006. eprint: https://www.jneurosci.org/content/26/16/4266.full.pdf.

[6] N. Burgess, C. Barry, and J. O’Keefe, “An oscillatory interference model of grid cell firing,” Hippocampus, vol. 17, no. 9, pp. 801–812, Jun. 2007. doi: 10.1002/hipo.20327.

[7] T. Waaga, H. Agmon, V. A. Normand, et al., “Grid-cell modules remain coordinated when neural activity is dissociated from external sensory cues,” Neuron, S0896627322002471, Apr. 2022. doi: 10.1016/j.neuron.2022.03.011.

[8] M. G. Campbell, S. A. Ocko, C. S. Mallory, et al., “Principles governing the integration of landmark and self-motion cues in entorhinal cortical codes for navigation,” Nature Neuroscience, vol. 21, no. 8, pp. 1096–1106, Aug. 2018. doi: 10.1038/s41593-018-0189-y.

[9] M. G. Campbell, A. Attinger, S. A. Ocko, et al., “Distance-tuned neurons drive specialized path integration calculations in medial entorhinal cortex,” Cell Reports, vol. 36, no. 10, p. 109–669, Sep. 2021. doi: 10.1016/j.celrep.2021.109669.

[10] G. Chen, Y. Lu, J. A. King, et al., “Differential influences of environment and self-motion on place and grid cell firing,” Nature Communications, vol. 10, no. 1, p. 630, Dec. 2019. doi: 10.1038/s41467-019-08550-1.

[11] S. S. Winter, M. L. Mehlman, B. J. Clark, et al., “Passive Transport Disrupts Grid Signals in the Parahippocampal Cortex,” Current Biology, vol. 25, no. 19, pp. 2493–2502, Oct. 2015. doi: 10.1016/j.cub.2015.08.034.

[12] C. Parron and E. Save, “Evidence for entorhinal and parietal cortices involvement in path integration in the rat,” Experimental Brain Research, vol. 159, no. 3, pp. 349–359, Dec. 2004. doi: 10.1007/s00221-004-1960-8.

[13] C. J. Cueva and X.-X. Wei, “Emergence of grid-like representations by training recurrent neural networks to perform spatial localization,” arXiv:1803.07770 [cs, q-bio, stat], Mar. 2018. arXiv: 1803.07770 [cs, q-bio, stat].

[14] B. Sorscher, G. C. Mel, S. A. Ocko, et al., “A unified theory for the computational and mechanistic origins of grid cells,” Neuron, Oct. 2022. doi: 10.1016/j.neuron.2022.10.003.

[15] A. Banino, C. Barry, B. Uria, et al., “Vector-based navigation using gridlike representations in artificial agents,” Nature, vol. 557, no. 7705, pp. 429–433, May 2018. doi: 10.1038/s41586-018-0102-6.

[16] R. Schaeffer, M. Khona, and I. R. Fiete, “No Free Lunch from Deep Learning in Neuroscience: A Case Study through Models of the Entorhinal-Hippocampal Circuit,” Neuroscience, Preprint, Aug. 2022. doi: 10.1101/2022.08.07.503109.

[17] R. Schaeffer, M. Khona, S. Koyejo, et al., Disentangling Fact from Grid Cell Fiction in Trained Deep Path Integrators, Dec. 2023. arXiv: 2312.03954 [q-bio].

[18] A. Nayebi, A. Attinger, M. G. Campbell, et al., “Explaining heterogeneity in medial entorhinal cortex with task-driven neural networks,” Neuro-science, Preprint, Nov. 2021. doi: 10.1101/2021.10.30.466617.

[19] V. Schøyen, M. B. Pettersen, K. Holzhausen, et al., “Coherently remap-ping toroidal cells but not Grid cells are responsible for path integration in virtual agents,” iScience, vol. 26, no. 11, p. 108–102, Nov. 2023. doi: 10.1016/j.isci.2023.108102.

[20] J. C. R. Whittington, W. Dorrell, S. Ganguli, et al., Disentangling with Biological Constraints: A Theory of Functional Cell Types, Sep. 2022. arXiv: 2210.01768 [cs, q-bio].

[21] D. Xu, R. Gao, W.-H. Zhang, et al., Conformal Isometry of Lie Group Representation in Recurrent Network of Grid Cells, Nov. 2022. arXiv: 2210.02684 [cs, q-bio, stat].

[22] R. Gao, J. Xie, X.-X. Wei, et al., On Path Integration of Grid Cells: Group Representation and Isotropic Scaling, Nov. 2021. arXiv: 2006.10259 [cs, q-bio, stat].

[23] R. Schaeffer, M. Khona, T. Ma, et al., Self-Supervised Learning of Representations for Space Generates Multi-Modular Grid Cells, Nov. 2023. arXiv: 2311.02316 [cs].

[24] W. Dorrell, P. E. Latham, T. E. J. Behrens, et al., Actionable Neural Representations: Grid Cells from Minimal Constraints, Feb. 2023. arXiv: 2209.15563 [q-bio].

[25] B. L. McNaughton, F. P. Battaglia, O. Jensen, et al., “Path integration and the neural basis of the ‘cognitive map’,” Nature Reviews Neuroscience, vol. 7, no. 8, pp. 663–678, Aug. 2006. doi: 10.1038/nrn1932.

[26] D. Bush, C. Barry, D. Manson, et al., “Using Grid Cells for Navigation,” Neuron, vol. 87, no. 3, pp. 507–520, Aug. 2015. doi: 10.1016/j.neuron.2015.07.006.

[27] S. Dang, Y. Wu, R. Yan, et al., “Why grid cells function as a metric for space,” Neural Networks, vol. 142, pp. 128–137, Oct. 2021. doi: 10.1016/j.neunet.2021.04.031.

[28] G. Ginosar, J. Aljadeff, L. Las, et al., “Are grid cells used for navigation? On local metrics, subjective spaces, and black holes,” Neuron, 2023. doi: 10.1016/j.neuron.2023.03.027.

[29] E. I. Moser and M.-B. Moser, “A metric for space,” Hippocampus, vol. 18, no. 12, pp. 1142–1156, 2008. doi: 10.1002/hipo.20483.

[30] I. R. Fiete, Y. Burak, and T. Brookings, “What Grid Cells Convey about Rat Location,” Journal of Neuroscience, vol. 28, no. 27, pp. 6858–6871, Jul. 2008. doi: 10.1523/JNEUROSCI.5684-07.2008.

[31] T. Solstad, E. I. Moser, and G. T. Einevoll, “From grid cells to place cells: A mathematical model,” Hippocampus, vol. 16, no. 12, pp. 1026–1031, Dec. 2006. doi: 10.1002/hipo.20244.

[32] H. Stensola, T. Stensola, T. Solstad, et al., “The entorhinal grid map is discretized,” Nature, vol. 492, no. 7427, pp. 72–78, Dec. 2012. doi: 10.1038/nature11649.

[33] M. Stemmler, A. Mathis, and A. V. M. Herz, “Connecting multiple spatial scales to decode the population activity of grid cells,” Science Advances, vol. 1, no. 11, e1500816, Dec. 2015. doi: 10.1126/science.1500816.

[34] K. Yoon, M. A. Buice, C. Barry, et al., “Specific evidence of low-dimensional continuous attractor dynamics in grid cells,” Nature Neuroscience, vol. 16, no. 8, pp. 1077–1084, Aug. 2013. doi: 10.1038/nn.3450.

[35] D. Wennberg, “The Distribution of Spatial Phases of Grid Cells,” p. 91,

[36] A. Guanella, D. Kiper, and P. Verschure, “A MODEL OF GRID CELLS BASED ON A TWISTED TORUS TOPOLOGY,” International Journal of Neural Systems, vol. 17, no. 04, pp. 231–240, Aug. 2007. doi: 10.1142/S0129065707001093.

[37] R. J. Gardner, E. Hermansen, M. Pachitariu, et al., “Toroidal topology of population activity in grid cells,” Nature, vol. 602, no. 7895, pp. 123–128, Feb. 2022. doi: 10.1038/s41586-021-04268-7.

[38] L. Kang, B. Xu, and D. Morozov, “Evaluating State Space Discovery by Persistent Cohomology in the Spatial Representation System,” Frontiers in Computational Neuroscience, vol. 15, p. 616–748, Apr. 2021. doi: 10.3389/fncom.2021.616748.

[39] D. Ganguli and E. P. Simoncelli, “Efficient sensory encoding and bayesian inference with heterogeneous neural populations,” Neural computation, vol. 26, no. 10, pp. 2103–2134, 2014.

[40] C. Curto, E. Gross, J. Jeffries, et al., “What makes a neural code convex?” SIAM Journal on Applied Algebra and Geometry, vol. 1, no. 1, pp. 222–238, 2017.

[41] C. Curto, E. Gross, J. Jeffries, et al., “Algebraic signatures of convex and non-convex codes,” Journal of pure and applied algebra, vol. 223, no. 9, pp. 3919–3940, 2019.

[42] M. Karoubi and C. Weibel, On the covering type of a space, Dec. 2016. arXiv: 1612.00532 [math].

[43] P. J. Heawood, “Map colour theorem,” Quarterly Journal of Mathematics, vol. 24, pp. 332–338, 1890.

[44] B. D. Ripley, “Modelling spatial patterns,” Journal of the Royal Statisti-cal Society: Series B (Methodological), vol. 39, no. 2, pp. 172–192, 1977. doi: 10.1111/j.2517-6161.1977.tb01615.x. eprint: https://rss.onlinelibrary.wiley.com/doi/pdf/10.1111/j.2517-6161.1977.tb01615.x.

[45] J. Besag, “Discussion on Dr Ripley’s Paper,” Journal of the Royal Statistical Society: Series B (Methodological), vol. 39, no. 2, pp. 192–212, Jan. 1977. doi: 10.1111/j.2517-6161.1977.tb01616.x.

[46] M. A. Kiskowski, J. F. Hancock, and A. K. Kenworthy, “On the Use of Ripley’s K-Function and Its Derivatives to Analyze Domain Size,” Biophysical Journal, vol. 97, no. 4, pp. 1095–1103, Aug. 2009. doi: 10.1016/j.bpj.2009.05.039.

[47] D. Derdikman and E. I. Moser, “A Manifold of Spatial Maps in the Brain,” in Space, Time and Number in the Brain, Elsevier, 2011, pp. 41–57. doi: 10.1016/B978-0-12-385948-8.00004-9.

[48] J. H. Wen, B. Sorscher, S. Ganguli, et al., “One-shot entorhinal maps enable flexible navigation in novel environments,” Neuroscience, Preprint, Sep. 2023. doi: 10.1101/2023.09.07.556744.

[49] C. N. Boccara, M. Nardin, F. Stella, et al., “The entorhinal cognitive map is attracted to goals,” Science, vol. 363, no. 6434, pp. 1443–1447, Mar. 2019. doi: 10.1126/science.aav4837.

[50] N. J. Killian, M. J. Jutras, and E. A. Buffalo, “A map of visual space in the primate entorhinal cortex,” Nature, vol. 491, no. 7426, pp. 761–764, 2012.

[51] D. Aronov, R. Nevers, and D. W. Tank, “Mapping of a non-spatial dimension by the hippocampal–entorhinal circuit,” Nature, vol. 543, no. 7647, pp. 719–722, Mar. 2017. doi: 10.1038/nature21692.

[52] R. B. Gabrielsson, B. J. Nelson, A. Dwaraknath, et al., “A topology layer for machine learning,” in International Conference on Artificial Intelligence and Statistics, PMLR, 2020, pp. 1553–1563.

[53] B. Rieck, N. Köhler, C. Bodnar, et al., Pytorch-topological, https://github.com/aidos-lab/pytorch-topological/tree/main, 2021.

[54] D. P. Kingma and J. Ba, “Adam: A Method for Stochastic Optimization,” arXiv:1412.6980 [cs], Jan. 2017. arXiv: 1412.6980 [cs].

[55] A. Paszke, S. Gross, F. Massa, et al., “PyTorch: An Imperative Style, High-Performance Deep Learning Library,” in Advances in Neural Information Processing Systems, vol. 32, Curran Associates, Inc., 2019.

[56] L. McInnes, J. Healy, and J. Melville, UMAP: Uniform Manifold Approximation and Projection for Dimension Reduction, Sep. 2020. arXiv: 1802.03426 [cs, stat].

[57] C. Tralie, N. Saul, and R. Bar-On, “Ripser.py: A lean persistent homology library for python,” The Journal of Open Source Software, vol. 3, no. 29, p. 925, Sep. 2018. doi: 10.21105/joss.00925.

[58] P. Virtanen, R. Gommers, T. E. Oliphant, et al., “SciPy 1.0: Fundamental Algorithms for Scientific Computing in Python,” Nature Methods, vol. 17, pp. 261–272, 2020. doi: 10.1038/s41592-019-0686-2.

